# Identification of key molecules and biological processes in TCF21 treated tumor pericytes

**DOI:** 10.1101/2022.05.02.490305

**Authors:** Guofang Zhao, Donghong Zhang, Mengshan Wang

**Affiliations:** Shanghai University of Medicine and Health Sciences, Shanghai, China

## Abstract

Colorectal cancer has become a major public health problem in the US. Transcription factor 21 (TCF21) is reported to be silenced in colorectal cancer tissues. However, the mechanism of TCF21 in tumor pericytes is still unclear. In our study, we aim to identify the key biological processes and signaling pathways by analyzing the RNA-seq data. The GSE200064 was produced by the Illumina NovaSeq 6000 (Homo sapiens). The KEGG and GO analyses showed that MAPK signaling pathway and complement/coagulation cascades are the major changed signaling pathways in the progression of tumor pericytes with overexpression of TCF21. Moreover, we identified several interactive molecules including VEGFA, MMP2, CCL2, COL3A1, COL1A2, CXCL12, ELN, PDGFRB, VWF, and APOE. These findings may benefit the study of colorectal cancer treatment.

## Introduction

Colorectal cancer is one of the most common cancers worldwide^1^. The median age at diagnosis is around 70 years in developed countries. Among them, the highest incidence is found in countries of Europe and North America, whereas incidence is lowest in central Asia and Africa^2^. Currently, the incidence has decreased in the USA, which is probably due to the increased use of sigmoidoscopy and colonoscopy^3^. Metastasis leads to most cancer death^4^. Unlike primary tumors that can be treated using local surgery or radiation, cancer metastasis is a global issue^5^. In metastatic cancer, for which checkpoint immunotherapy was less effective. Thus, curing cancer metastasis remains a challenge^6^. Though cancer cell metastasis can start early time, most cells cannot colonize distant organs^7^. To form metastasis, cancer cells will undergo a series of steps. Pericytes are related to stabilizing blood vessel structure and permeability in cancer^8^. Pericytes affect tumor metastasis both positively and negatively. Most tumors favor establishing a stable vascular network to nourish tumor cells by maintaining vascular permeability. These aspects require a number of pericytes and depletion of pericytes results in tumor regression^9^.

In this study, we analyzed tumor pericytes with the overexpression of TCF21 by using the RNA-seq data. We determined several DEGs and significant signaling pathways.

We then performed the gene enrichment and created protein-protein interaction (PPI) network to understand the interacting relationships. These genes and biological processes may provide insights into the treatment of colorectal cancer.

## Methods

### Data resources

Gene dataset GSE200064 was obtained from the GEO database (https://www.ncbi.nlm.nih.gov/geo/). The data was created by the Illumina NovaSeq 6000 (Homo sapiens) (Jinan University, HuangPu street, GuangZhou 510632, China). The analyzed dataset includes three groups of tumor pericytes and three groups of tumor pericytes with the overexpression of TCF21.

### Data acquisition and processing

The data were organized and conducted by the R package as previously described ^10–18^. We used a classical t-test to identify DEGs with *P* < 0.05 and fold change ≥ 1.5 as being statistically significant.

The Kyoto Encyclopedia of Genes and Genomes (KEGG) and Gene Ontology (GO) KEGG and GO analyses were performed by the R package (ClusterProfiler) and Reactome (https://reactome.org/). *P* < 0.05 was considered statistically significant.

### Protein-protein interaction (PPI) networks

The Molecular Complex Detection (MCODE) was used to construct the PPI networks^19–24^ The significant modules were produced from PPI networks and String networks (https://string-db.org/). The biological processes analyses were performed by using Reactome, and *P* < 0.05 was considered significant.

## Results

### Identification of DEGs in tumor pericytes with TCF21 overexpression

To determine the effects of TCF21 on tumor pericytes from patients with colorectal cancer, we analyzed the RNA-seq data of tumor pericytes with the overexpression of TCF21. A total of 827 genes were identified with a threshold of *P* < 0.05. The top up- and down-regulated genes were shown by the heatmap and volcano plot (Figure 1). The top ten DEGs were listed in Table 1.

**Figure 1.**
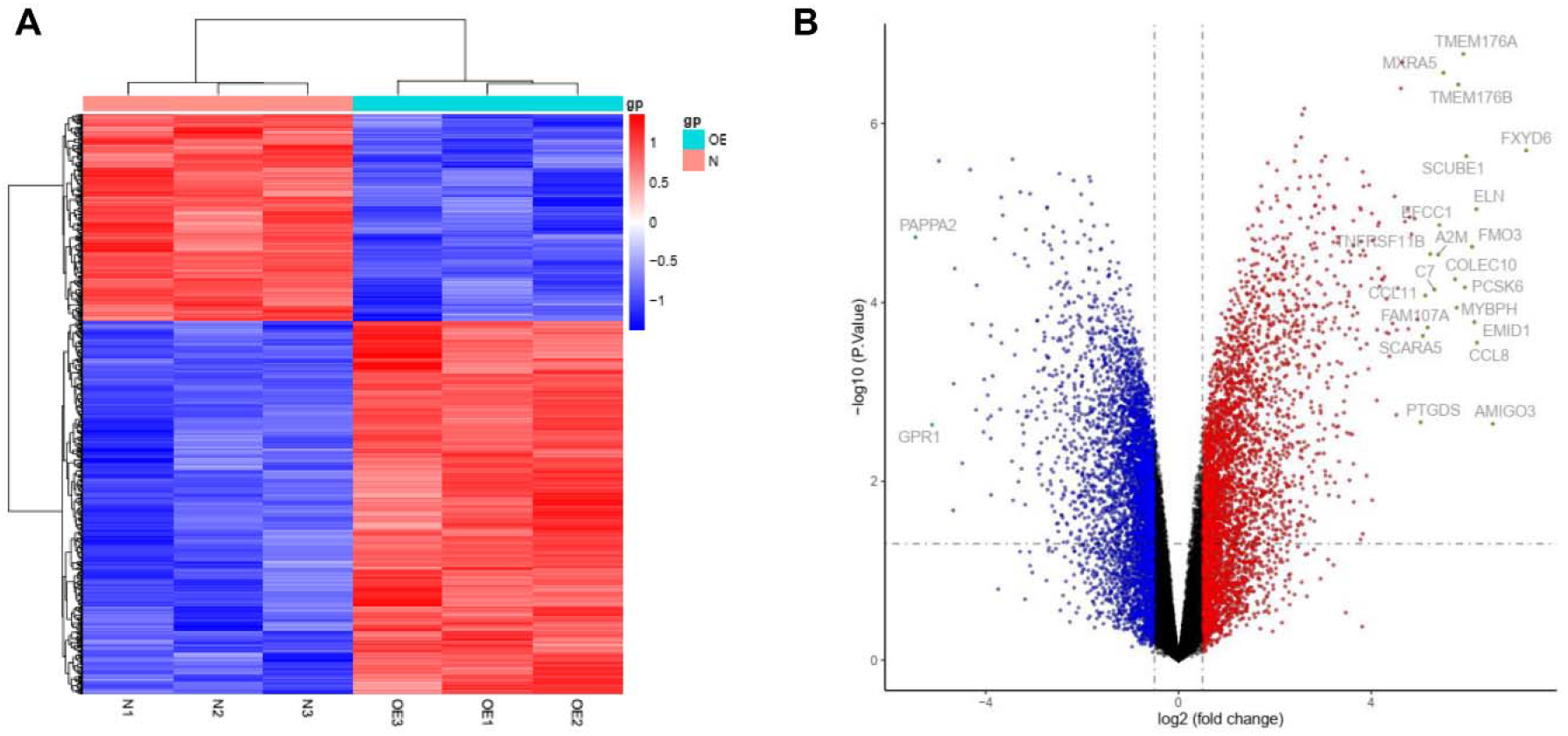
Heatmap and volcano plot of DEGs in tumor pericytes with TCF21 overexpression. (A) Significant DEGs (*P* < 0.05) were used to create the heatmap. (B) Volcano plot for DEGs plot in tumor pericytes with TCF21 overexpression. Note that the most significantly changed genes are highlighted by grey dots.

**Table 1.**
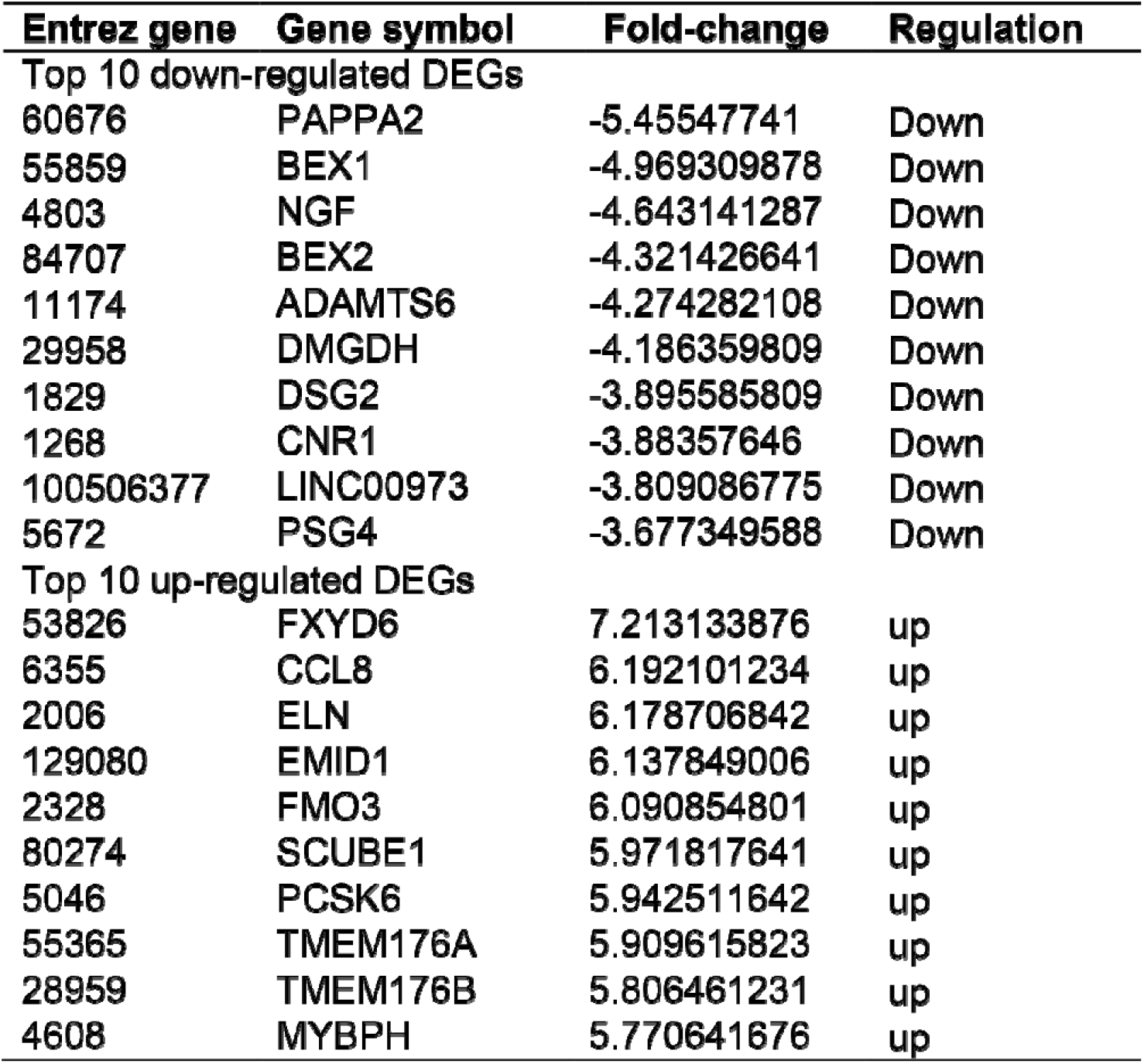

**Table 2.**
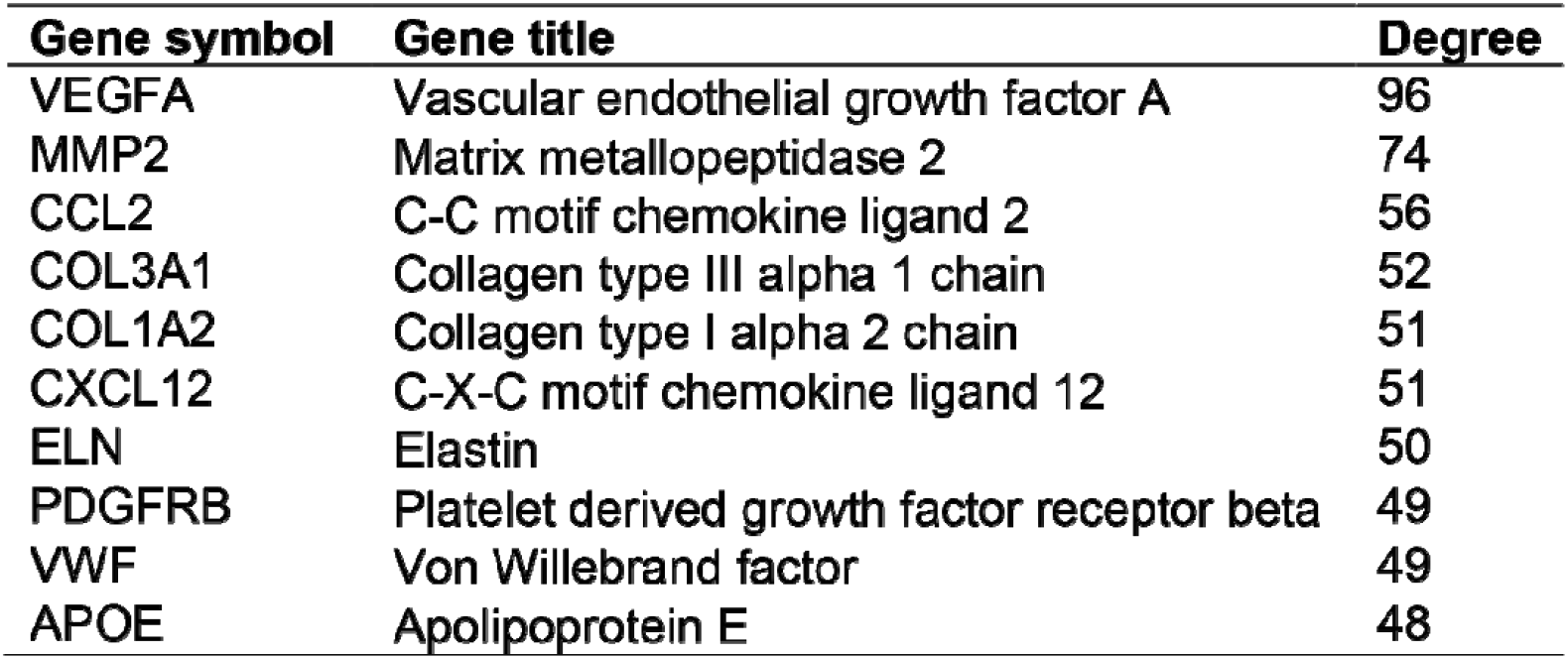
Top ten genes demonstrated by connectivity degree in the PPI network.

### Enrichment analysis of DEGs in tumor pericytes with TCF21 overexpression

To identify the mechanism of TCF21 in tumor pericytes, we analyzed the enrichment of KEGG and GO items (Figure 2). We identified the top ten KEGG items, including “MAPK signaling pathway”, “Complement and coagulation cascades”, “Protein digestion and absorption”, “Cell adhesion molecules”, “AGE-RAGE signaling pathway in diabetic complications”, “Staphylococcus aureus infection”, “Pertussis”, “TGF-beta signaling pathway”, “Steroid biosynthesis”, and “Terpenoid backbone biosynthesis”. We also identified the top ten biological processes (BP) of GO, including “extracellular matrix organization”, “extracellular structure organization”, “external encapsulating structure organization”, “cell-substrate adhesion”, “transmembrane receptor protein serine/threonine kinase signaling pathway”, “steroid metabolic process”, “urogenital system development”, “kidney development”, “renal system development”, and “sterol biosynthetic process”. We identified the top ten cellular components (CC) of GO, including “collagen-containing extracellular matrix”, “cell-cell junction”, “focal adhesion”, “cell-substrate junction”, “endoplasmic reticulum lumen”, “collagen trimer”, “basement membrane”, “platelet alpha granule”, “platelet alpha granule lumen”, and “complex of collagen trimers”. We identified the top ten molecular functions (MF) of GO, including “extracellular matrix structural constituent”, “glycosaminoglycan binding”, “sulfur compound binding”, “growth factor binding”, “heparin binding”, “integrin binding”, “collagen binding”, “extracellular matrix structural constituent conferring tensile strength”, “extracellular matrix binding”, and “platelet-derived growth factor binding”.

**Figure 2.**
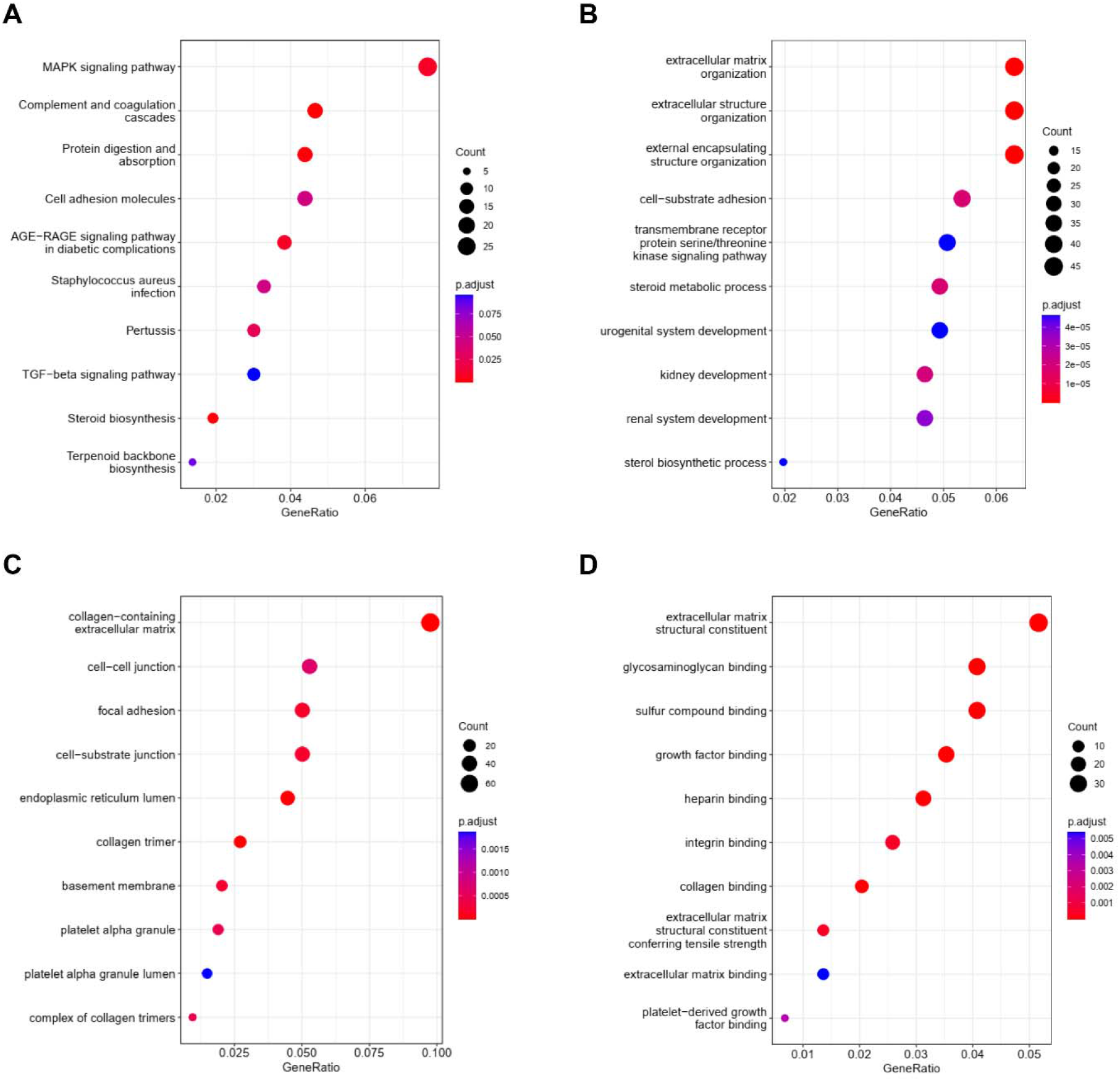
KEGG and GO analyses of DEGs in tumor pericytes with TCF21 overexpression. (A) KEGG: Kyoto Encyclopedia of Genes and Genomes, (B) BP: Biological processes, (C) CC: Cellular components, (D) MF: Molecular functions.

### PPI network and Reactome analyses

To explore the potential interactions among the DEGs, we created the PPI network by using 789 nodes and 2693 edges. The combined score > 0.2 was set as a cutoff by using the Cytoscape software. Table 2 showed the top ten genes with the highest scores. The top two significant modules were shown in Figure 3. We further analyzed the PPI and DEGs with a Reactome map (Figure 4) and identified the top ten biological processes including “Extracellular matrix organization”, “Attenuation phase”, “Activation of gene expression by SREBF (SREBP)”, “HSF1-dependent transactivation “, “Assembly of collagen fibrils and other multimeric structures”, “NGF-stimulated transcription”, “Regulation of cholesterol biosynthesis by SREBP (SREBF)”, “Degradation of the extracellular matrix”, “Collagen formation”, “Nuclear Events (kinase and transcription factor activation)” (Supplemental Table S1).

**Figure 3.**
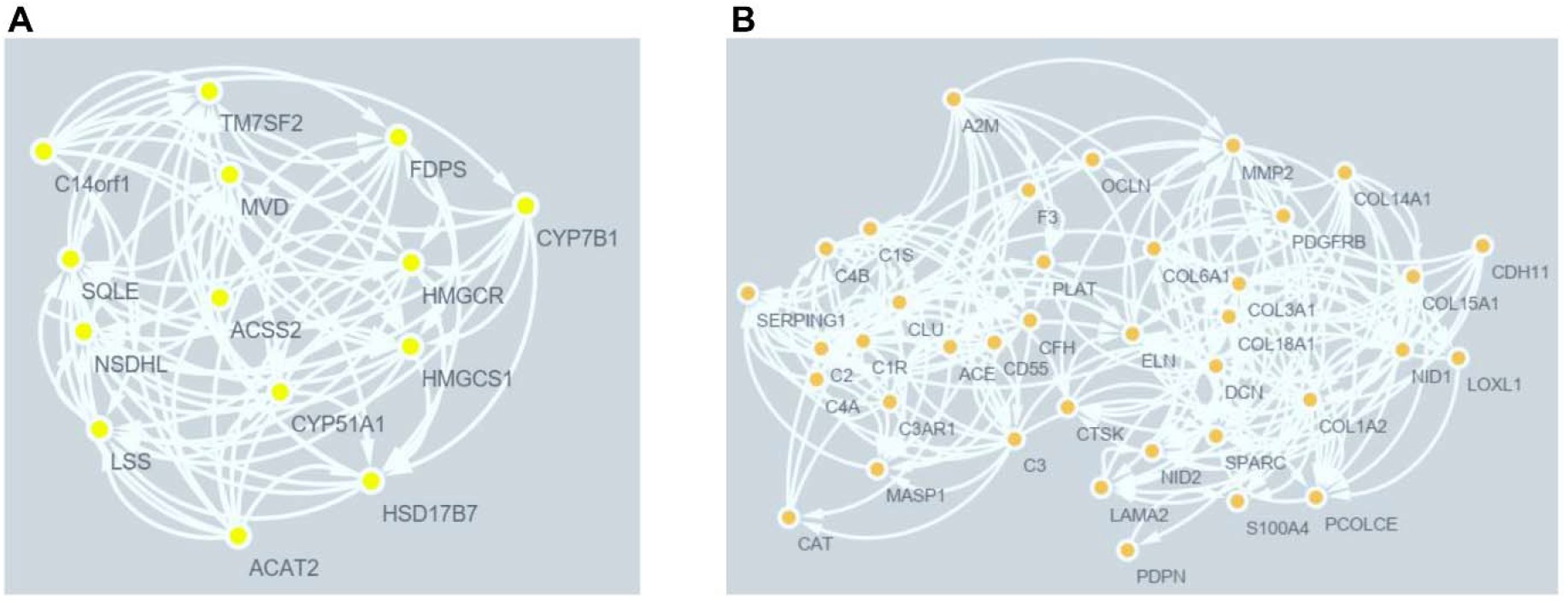
The PPI network analyses of DEGs in tumor pericytes with TCF21 overexpression. The cluster (A) and cluster (B) were created by MCODE by using the Cytoscape.

**Figure 4.**
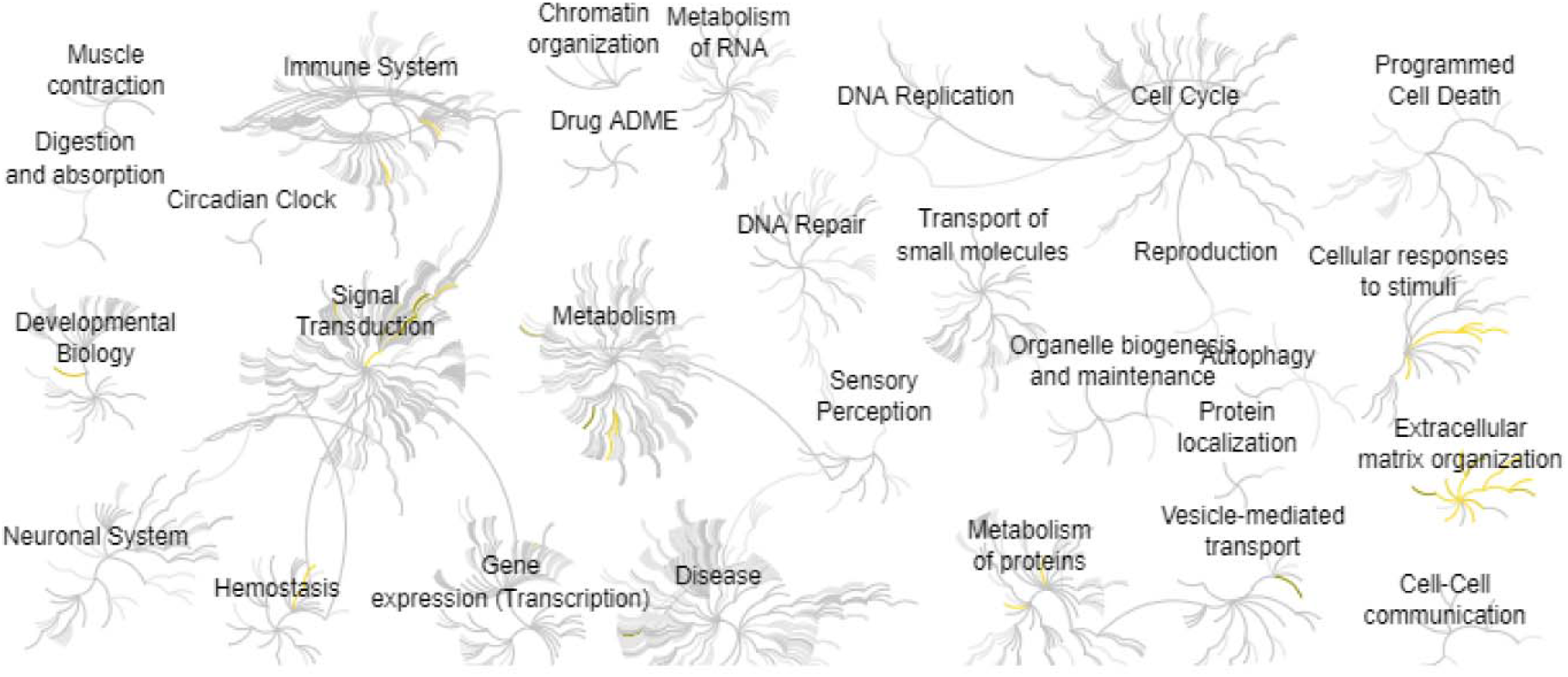
Reactome map representation of the significant biological processes in tumor pericytes with TCF21 overexpression.

## Discussion

Transcription factor 21 (TCF21) is one of the basic helix-loop-helix (bHLH) transcription factors that control cell fate and differentiation that plays a critical role in a wide spectrum of biological processes, including organogenesis, proliferation, differentiation as well as invasion and metastasis of cancer cells^25^. Controlled expression and activity of TCF21 provide a changed transcriptional program that affects the progression of cancer^26^.

In our study, we found that the “MAPK signaling pathway” and “Complement and coagulation cascades” are the major changed signaling pathways in tumor pericytes with the overexpression of TCF21. The MAPK pathways are downstream of many growth-factor receptors and overexpression of these receptors is commonly detected in colorectal cancer^27^. The circadian clock is a key regulator for a number of pathological processes including aging, inflammation, metabolism, and cancer^28–38^. Interestingly, MAPKs have important roles in the regulation of the circadian biological clock. In the mammalian suprachiasmatic nucleus (SCN), MAPK pathways can function as inputs to 24h cycles that are consistent with the circadian clock^39^. Jing Zhang et al found that complement and coagulation cascades signaling is related to the soft tissue sarcoma^40^.

By creating the PPI network, we also identify the ten key molecules in colorectal cancer cells with the overexpression of TCF21. Haihong Pu et al found that VEGFA is closely associated with breast cancer^41^. Liping Han et al found that MMP2 expression is related to lung cancer and clinical parameters^42^. Su Yin Lim et al found that CCL2-CCR2 signaling is a key process in cancer metastasis^43^. G protein-coupled receptor (GPCR) and regulators of G protein signaling (RGS) family proteins are involved in a variety of diseases such as cancer, arthritis, metabolic dysfunctions, and aging-related diseases^44–56^ Wendy Kleibeuker et al found that physiological changes of G protein-coupled receptor kinase 2 (GRK2) control CCL2-induced signaling^57^. Lushun Yuan et al found the overexpression of COL3A1 leads to a poor prognosis in bladder cancer^58^. Yifan Yu et al found there are inhibitory effects of COL1A2 on colorectal cancer cell proliferation and migration^59^. Weiqiang Zhou et al found targeting the CXCL12 is of critical importance for tumor immunotherapy^60^. Laura Jamrog et al found that PAX5-ELN leads to multistep B-cell leukemia in mice^61^. Yingchi Zhang et al found that PDGFRB mutation is a characteristic of Ph-like acute lymphoblastic leukemia^62^. Lubor Borsig et al found VWF results in thrombosis during cancer^63^.

To sum up, this study found the significant molecules and signaling pathways during the overexpression of TCF21 in tumor pericytes. The “MAPK signaling pathway” and “Complement and coagulation cascades” are the major changed signaling pathways. Our study may provide novel ideas for the treatment of colorectal cancer.

## Author Contributions

Guofang Zhao, Donghong Zhang: Methodology and Writing. Mengshan Wang: Conceptualization, Writing-Reviewing and Editing.

